# Pentosan polysulfate inhibits attachment and infection by SARS-CoV-2 *in vitro*: insights into structural requirements for binding

**DOI:** 10.1101/2021.12.19.473359

**Authors:** Sabrina Bertini, Anna Alekseeva, Stefano Elli, Isabel Pagani, Serena Zanzoni, Giorgio Eisele, Ravi Krishnan, Klaus P Maag, Christian Reiter, Dominik Lenhart, Rudolf Gruber, Edwin A Yates, Elisa Vicenzi, Annamaria Naggi, Antonella Bisio, Marco Guerrini

## Abstract

Two years since the outbreak of the novel coronavirus SARS-CoV-2 pandemic, there remain few clinically effective drugs to complement vaccines. One is the anticoagulant, heparin, which in 2004 was found able to inhibit invasion of SARS CoV (CoV-1) and which has been employed during the current pandemic to prevent thromboembolic complications and moderate potentially damaging inflammation. Heparin has also been shown experimentally to inhibit SARS-CoV-2 attachment and infection in susceptible cells. At high therapeutic doses however, heparin increases the risk of bleeding and prolonged use can cause heparin-induced thrombocytopenia, a serious side-effect. One alternative, with structural similarities to heparin is the plant-derived, semi-synthetic polysaccharide, pentosan polysulfate (PPS). PPS is an established drug for the oral treatment of interstitial cystitis, is well-tolerated and exhibits weaker anticoagulant effects than heparin. In an established Vero cell model, PPS and its fractions of varying molecular weights, inhibited invasion by SARS-CoV-2. Intact PPS and its size-defined fractions were characterized by molecular weight distribution and chemical structure using NMR spectroscopy and LC-MS, then employed to explore the structural basis of interactions with SARS-CoV-2 spike protein receptor-binding domain (S1 RBD) and the inhibition of Vero cell invasion. PPS was as effective as unfractionated heparin, but more effective at inhibiting cell infection than low molecular weight heparin (on a weight/volume basis). Isothermal titration calorimetry and viral plaque-forming assays demonstrated size-dependent binding to S1 RBD and inhibition of Vero cell invasion, suggesting the potential application of PPS as a novel inhibitor of SARS-CoV-2 infection.

## Introduction

The enveloped, positive sense RNA virus, SARS-CoV-2, belonging to the *Coronaviridae*, is responsible for the COVID-19 pandemic and causes serious clinical morbidities that are consistent with the onset of severe acute respiratory distress syndrome (SARS).^1^ By late 2021, the total number of COVID-19 deaths worldwide stood at over 5 million, but this is probably an underestimate owing to the difficulties of collecting accurate data globally. Despite the significant contribution of vaccines to preventing much severe morbidity and mortality caused by SARS-CoV-2 infection, there remains an urgent need to develop additional drugs to treat or prevent the infection. COVID-19 is a disease which continues to occur due to the waning of immunity, the infection of immunosuppressed patients or those with underlying illness, as well as the emergence of mutations and the resulting breakthrough infections. Several drugs, marketed for other therapeutic indications, have been re-purposed to treat COVID-19 patients and antiviral strategies that include treatment with remdesivir or convalescent plasma, have received emergency approval.^2,3^ Despite promising results, the use of such treatments is limited, owing to their expense and, because they can only be delivered intravenously. Additional treatments are therefore required and, indeed, during the writing of this article, the first orally available antiviral drug against COVID, molnupiravir, was approved for use in the UK.^4,5^

One existing family of drugs which have been re-purposed for treatment of COVID-19, are heparins. Heparins are a class of polydisperse, linear, sulfated polysaccharides of animal origin. Besides their well-known anticoagulant properties, heparins have been shown to inhibit cell invasion by SARS CoV-2^6,7^, SARS CoV-1^8,9^, influenza H5N1^10^, as well as several flaviviruses, including Dengue and Yellow Fever viruses.^11^ Heparin can also influence the ability of the infected cell to survive, as evinced by a study of the survival of cells infected by Zika virus^12^. The activity of heparin against SARS-CoV-2, has been established using a number of *in vitro* experimental models and its efficacy appears to depend largely in its interactions with the spike protein 1 receptor binding domain (S1-RBD). ^6,7,13^

Regarding the clinical use of heparin, a systematic review and meta-analysis of the association of anticoagulant status and mortality in COVID-19 patients was conducted encompassing 29 articles that had been published to January 2021. The study concluded that both therapeutic and prophylactic anticoagulant regimes reduced in-hospital mortality compared with untreated patients.^14^ Anticoagulant therapy has been recommended in the prophylaxis of thromboembolism in COVID patients with respiratory infection and reduced mobility.^15^ Nevertheless, the occurrence of bleeding complications and the documented risk of heparin–induced thrombocytopenia (HIT) after prolonged exposure, both limit the use of heparin in patients having high incidence of VTE, particularly for high dose administration.

The proposed mechanism of inhibition by heparin involves competition for the binding of the SARS-CoV-2 spike protein (S1) with endogenous cell surface heparan sulfate (HS), a co-receptor which serves as an anchor and to facilitate the conformational change that enables viral binding to angiotensin-converting enzyme 2 (ACE2) ^6,7^ The high negative charge density of heparin, a close structural analogue of HS, favours interaction with the positively charged amino acid residues of S1 (R346, R355, N354, R357, K444 and R466).^6,16^ The results from several studies have suggested that other sulfated molecules and highly charged polyelectrolytes could also provide potential targets for the design of inhibitors against the entry of viruses.^17,18^

Pentosan polysulfate (PPS) is a chemically-sulfated derivative of pentosan, a plant-derived xylan that has been approved for the treatment of bladder pain and discomfort in interstitial cystitis (ema.europa.eu/en/medicines/human/EPAR/elmiron) and could represent an alternative to heparin. PPS is produced by the exhaustive chemical O-sulfation of hardwood glucuronoxylan whose backbone comprises a linear polymer of β-D-xylopyranose (Xyl) units linked β (1–4) to each other and branched with 4-O-methyl-glucuronic acid (MGA) residues linked α (1-2) to the xylose backbone^19^ (**Fig. 1a**). PPS has poor anticoagulant activity and is known for its anti-inflammatory activities and affinity for the cytokines, IL-4, IL-5 and IL-13, resulting in similar anti-inflammatory efficacy as topical steroids in a guinea pig model of allergic rhinitis.^20^

**Fig. 1.**
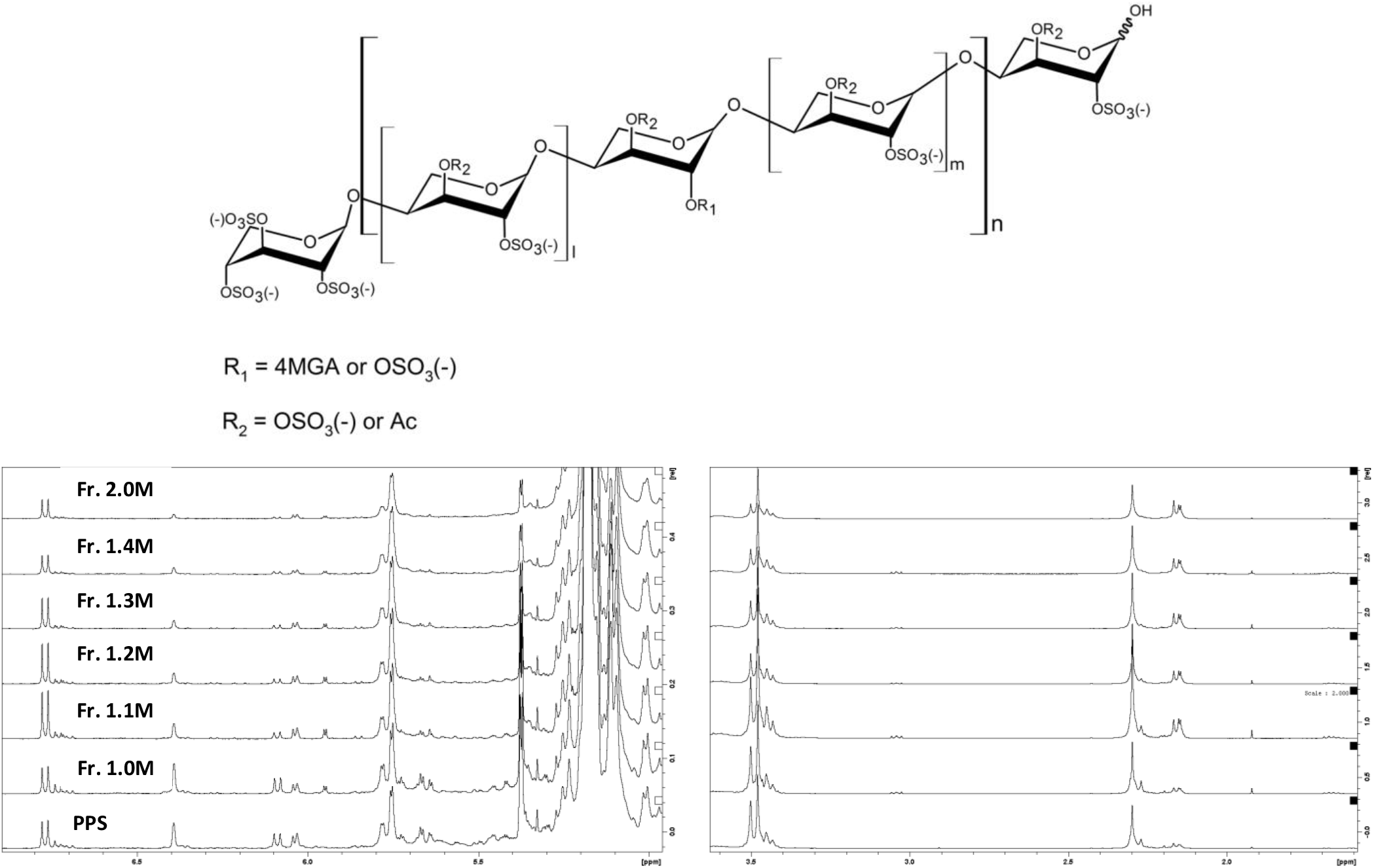
(A) Major structural signatures of PPS. MGA – branching 4-O-methyl-2,3-di-O-sulfated glucuronic acid, Ac – acetyl group (B) Low field and high field proton spectral regions of PPS and its fractions.

In the present study, a sample of PPS (API) was fractionated according to its charge density and molecular dimension using anion exchange chromatography. The parent polysaccharide and six fractions were characterized in terms of their molecular weight distribution, as well as by NMR spectroscopy and LC-MS analysis. These were all then evaluated for their ability to bind the S1-RBD and to inhibit virus invasion in comparison with both unfractionated heparin (UFH) and its low molecular weight heparin counterpart (LMWH). The reduced anticoagulant potency of PPS and its fractions compared to heparin, alongside its well-tolerated oral administration even at high dose (300 mg daily)^21^ suggest that oral administration of PPS might provide an effective and safe prophylactic treatment of COVID-19 disease.

## Methods & Materials

### Preparation of PPS fractions

Native PPS was fractionated on a DEAE Sephacel™ column applying aqueous NaCl (0.6M – 2.0M, stepwise elution) as eluent. PPS fractions were precipitated by addition of a twofold excess of ethanol and storing at −30 °C over night. Samples were collected by centrifugation (3500 rpm, 10 min), washed with 70% ethanol and dried in vacuo at 40 °C.

### NMR characterization of PPS and PPS fractions

^1^H-NMR and HSQC NMR spectra were acquired using a Bruker Avance 500 NEO instrument (Bruker, Karlsruhe, Germany), equipped with 5 mm cryoprobe. *Circa* 40 mg of samples were weighed and dissolved in 0.5 ml of deuterated phosphate buffer solution, 0.15 mM, pH 7.1. Proton spectra were recorded at 303K, with pre-saturation of solvent during relaxation delay. Recording and integration of quantitative HSQC spectra, as well as quantification of the monosaccharide building blocks were performed according to the published study.^19^

### Molecular weight of PPS and PPS fractions

The molecular weight distribution of PPS and PPS-derived fractions were performed on a Viscotek 305 HPLC system (Viscotek, Houston, USA) equipped with a triple detector array exploiting simultaneous action of refraction index (RI) detector, viscometer and right-angle laser light-scattering (RALLS) detector, employing an adaptation of the published method.^22,23^ The applied RI increment (dn/dc) was found to be 0.093 ml/g and was used to convert the RI response to concentration.^19^

### LC-MS characterization of PPS and PPS fractions

Analyses by LC-MS of unfractionated PPS and its fractions were performed by ion pair reversed phase chromatography coupled with high resolution mass spectrometry as previously described.^19^ Briefly, the separation of PPS oligomers was achieved on a HPLC 1100 (Agilent, Santa Clara, USA) system equipped with a high resolution ESI-FTICR mass spectrometer Solarix 7T (Bruker Daltonics, Billerica, MA, USA), using a Kinetex 2.6 μm C18 column (100 × 2.1mm I.D., Phenomenex, Torrance, CA, USA) and dibutylamine acetate as ion pairing agent (eluent A and B containing 10 mM DBA and 10 mM acetic acid in water and methanol, respectively). All reagents were of LC-MS grade. The separation was achieved using a multi-step gradient at 0.1 ml/min flow-rate: the solvent composition was held at 35% B for the first 3 min, then increased to 55% B over 2 min, and to 75% B over another 35 min, where it was held at 75% for 20 min; afterwards, the content of the eluent B was increased to 98% in 30 min, where it was held for 10 min in order to elute the longest components. Finally, it was returned to 35% B over 2 min, and held for the last 23 min for equilibrating the chromatographic column before the next run. The following ESI-MS parameters were used: capillary voltage, 3.2 kV; nebulizer gas pressure, 1 bar; drying gas, 3.7 ml/min and 180°C. The mass spectra acquisition was performed in m/z 200-3000 range with time domain 1M and ion source accumulation of 300 ms. Injection volume was 3 μl for both 5 mg/ml unfractionated PPS and 2.5 mg/ml fractions. The fraction, Fr.2.0M, was not analysed by LC-MS due to poor chromatographic separation and the poor ionization of high molecular weight oligosaccharides, which results in low MS sensitivity.

### Dynamic light scattering

Dynamic light scattering (DLS) experiments were performed using a Zetasizer Nano ZS instrument (Malvern Panalytical, Malvern, UK) operating at 633 nm with backscatter detection at 173°. The protein solution (2 μM) was allowed to equilibrate with or without the addition of salt at the measurement temperature of 25 °C before starting acquisition. To analyze the S1-RBD/PPS complex size, the solutions were obtained by mixing S1-RBD with unfractionated and fractionated and PPS solutions in 20 mM HEPES buffer with salt at different protein ligand molar ratios. The complexes were transferred to a disposable cell at room temperature and analyzed 2 minutes post-mixing. Data were analyzed using Zetasizer software version 7.11 (Malvern Panalytical, Malvern, UK) and each value corresponds to the average of three separate measurements.

### Circular Dichroism

Circular dichroism (CD) spectra data were collected at 25°C on a Jasco J-1500 instrument (JASCO, Great Dunmow, UK) equipped with a Peltier unit-controlled cell holder. The spectra in the far-UV region (200–250 nm) were recorded using a quartz cell (Hellma, Southend-on-Sea, UK) with a path length of 0.1 cm, a response time of 1 s, a scan speed of 20 nm/min and bandwidth of 0.5 nm and each spectrum was the average of five scans. To address the conformational state of the S1-RBD domain in HEPES buffer, CD data were acquired on S1-RBD (4-10 μM) samples just after the reconstituted with de-ionised water as a control and after buffer exchange (HEPES buffer, pH 7.4 with NaCl 200mM). To analyse the conformational changes upon PPS complex formation, the S1-RBD solutions were incubated with unfractionated and fractionated PPS solutions in HEPES buffer at different protein ligand molar ratios and CD spectra were recorded immediately.

### Isothermal titration calorimetry

Isothermal titration calorimetry was used to characterize the binding affinities between PPS fractions and the S1-RBD domain. The experiments were carried out at 25 °C using a MicroCal PEAQ-ITC (Malvern Ltd., Malvern, UK). Both protein and ligand were prepared in PBS or HEPES buffer pH 7.2. Briefly, SARS-CoV-2-RBD protein (Sino Biological China) was firstly reconstituted with MilliQ filtered water and then, to remove the excess of trehalose, the buffer was exchange with PBS or 20 mM HEPES pH 7.2, NaCl 60 mM using an Amicon Ultra concentrator (10 kDa filter, 0.5 ml). Each lyophilized PPS fraction was initially resuspended with 500 ul of the same buffer. The protein and ligand stock solutions were than diluted to different concentrations for ITC analysis with PPS or HEPES buffer and 140 or 200 mM salt. For all the experiments S1-RBD was taken in the cell (2-3 μM) and PPS fractions in the syringe (30 μM). Each ITC experiment consisted of twenty injections of 2 μl each with an initial delay of 180 s, 180 s between injections, and a stirring rate of 500 rpm. To estimate the thermodynamic parameters (K_D_, ΔH and ΔS), the data were fitted using the MicroCal analysis software.

### Vero cell culture and assays

Vero cells were plated at 2.5×10^5^ cell/well in 24-well plates in EMEM supplemented with 10% fetal bovine serum (complete medium). Twenty-four hours later, three different PPS protocol treatments were adopted:

i. *Cell Treatment*- Cells were incubated with serial dilutions of PPS in 250 μl of complete medium for 30 min and then 50 μl of supernatant containing 50 plaque forming units (PFU) of SARS-CoV-2 isolate GISAID accession ID: EPI_ISL_413489 were added. ^6^
ii. *Virus Treatment*- Fifty plaque forming units (PFU) of SARS-CoV-2 in 50 μl of complete medium were incubated with compound serial dilutions of PPS for 30 min at 37 °C and then added to Vero cells in a final volume of 300 μl.
iii. *Cell + Virus Treatment*:- Fifty PFU of SARS-CoV-2 in 50 μl of complete medium were incubated with compound serial dilutions and cells were incubated with serial dilutions of PPS within the same range for 30 min at 37 °C in 250 μl. After 30 min of incubation at 37 °C, the virus containing supernatant was added to the PPS-treated cells.

In all three PPS protocol treatments, the incubation was then extended for 1 h at 37°C. Then, supernatants were discarded and, 500 μl of 1% methylcellulose overlay dissolved in medium containing 1% of fetal bovine serum were then added to each well. After 3 days, cells were fixed using a 6% (v/v) formaldehyde:phosphate-buffered saline solution and stained with 1% (w/v) crystal violet (Sigma-Aldrich, Italy) in 70% (v/v) methanol (Sigma-Aldrich, Italy). The plaques were counted under a stereoscopic microscope (SMZ-1500, Nikon).

### Statistical Analysis

Prism GraphPad software v. 9.0 (www.graphpad.com) was used for the statistical analyses. Comparison among groups were performed using the one-way analysis of variance (ANOVA). The mean of each treatment column was compared with the control column (Nil; no treatment) applying the Bonferroni correction. **** indicates a *p* value <0.0001, *** indicates a *p* value <0.001 and ** indicates a *p* value <0.01.

### Coagulation assays

All fractions were stored frozen (−20°C) at 1 mg/ml in Aqua-Dest. Dilutions in human pool plasma were used with the following concentrations (first dilution step to 100 μg/ml in 0.9 % NaCl) (n = 6): 0; 3.125; 6.25; 12.5; 25 and 50 μg/ml. The activated partial thromboplasin time (aPTT; SyntASil TM; reference range: 25 - 38 sec) and anti-Xa activity (Liquid anti-Xa TM; reference range 0 IU/ml) were measured with commercially available assays according to the manufacturer’s instructions (Werfen, Munich, Germany) and were analyzed on the ACL TOP analyzer (Werfen; Munich, Germany).

### PPS structural optimization and molecular docking simulations of PPS-SARS-CoV-2 RBD domain

A model comprising the linear hexasaccharide portion of PPS was represented by the xylohexasaccharide: 2,3,4-trisulfo-Xyl-β(1-4)-[2,3-disulfo-Xyl]_5_-OH. Two limit conformations of this xylohexasaccharide, characterized by all 2,3-disulfo-Xyl residues in ^1^C_4_ and ^4^C_1_ conformations, were built. In each 2,3-disulfo-Xyl residue, the conformation of the sulfate groups was manually adjusted to reduce the steric hindrance considering their orientation as axial or equatorial corresponding to the ^1^C_4_ and ^4^C_1_ chairs. In both models, the φi/ψi glycosidic backbone conformation was also manually adjusted before the final energy minimization step, to reduce the steric hindrance of the glycan chain. The final conformations of these two xylohexasaccharides having Xyl residues in ^1^C_4_ and ^4^C_1_ were energy minimized using the Maestro 12.7 Macromodel 13.1 software and molecular mechanic Amber* force-field (Suite 2021-1, Schrödinger Inc. San Diego) (**Fig. S5**). The non-bonded distance cut off were set as 20.0, 8.0, and 4.0 Å for electrostatic, Van der Waals and hydrogen bond potential energy interactions, respectively. The minimization procedure involved 10000 steps (default energy minimization algorithm) or a gradient threshold 1.0×10-3 KJ mol-1 Å-1, before the docking stage. Only the xylohexasaccharides in ^1^C_4_ (see result section) was submitted to molecular docking.

The hexasaccharide was docked on the surface of the S1-RBD (PDB ID 6M0J) targeting the sequence of positively charged patches R346, R355, K356, R357, that present good accessibility when the S1-RBD is in the ‘up’ conformation in the trimeric protein S (PDB ID 6VSB). All the hexasaccharide torsional degrees of freedom were adjusted during automatic docking, except for the glycosidic dihedrals φ_i_/ψ_i_ (_i_ =1, 5), while the S1-RBD-S1 was set as rigid. The docking grid was set as L_x_ = L_y_ = 80, L_z_ = 120 points, while the center of the grid was: (CM_x_, CM_y_, CM_z_) = (−40.272, 25.851, 21.266). The grid spacing was 0.375 Å (default). The software Autodock 4.2 was used for the automatic docking. The Lamarckian genetic algorithm 4.2 was used for the docking in combination with the following set of parameters: number of runs = 100, population size = 2000, maximum number of energy evaluations = 2.5 x 10^7^, maximum number of generations = 2.7 x 10^5^. After the search procedure, the obtained poses of the ligand-receptor complex were clustered (tolerance RMSD < 2.0 Å) and each cluster characterized by the lowest binding energy (autodock score function) and highest population (number of poses for cluster), were selected.

## Results

### Isolation and structural characterization of PPS fractions

To correlate the structural properties of PPS with their ability to bind the spike protein and inhibit virus propagation, PPS was fractionated by anion exchange chromatography and fractions were characterized by light scattering detection, NMR and LC-MS.

The molecular weights of PPS and its fractions were obtained by size exclusion chromatography coupled with a triple detector array, consisting of light scattering, refractive index and viscometer, following a similar procedure to that adopted for unfractionated heparin and low molecular weight heparins.^22,23^ The advantage of this method compared with classical approaches is that it does not require column calibration with reference standards possessing the same chemical structure as the samples to be analysed and exhibiting well defined molecular weight and narrow polydispersity. The accuracy of such relative measurements is therefore affected by the quality of the standards employed. The distinct elution times of PPS and its fractions reflects their different molecular weight and molecular weight distribution (**Fig. S1**). The M_w_ (weight average molecular weight), M_n_ (number average molecular weight), polydispersity D (expressed as M_w_/M_n_), Rh and η (dl/g), values obtained for the samples are reported in **Table 1**. The Mw values of the fractions ranged between 3.7 and 17.3 kDa, corresponding to an average chain length of 11 to 50 monosaccharide units.

**Table 1.** Properties of PPS and its fractions. The results refer to the mean values of duplicate injections. *Hydrodynamic radius; **intrinsic viscosity

| Sample | Mw (kDa) | Mn (kDa) | D (Mw/Mn) | Rh* | $\eta^{**}$ (dl/g) |
| --- | --- | --- | --- | --- | --- |
| (PPS) | 5.7 | 4.4 | 1.3 | 1.6 | 0.01 |
| (Fr. 1.0M) | 3.7 | 3.6 | 1.0 | 1.3 | 0.04 |
| (Fr. 1.1M) | 5.3 | 5.3 | 1.0 | 1.6 | 0.05 |
| (Fr. 1.2M) | 7.0 | 6.9 | 1.0 | 1.9 | 0.06 |
| (Fr. 1.3M) | 9.2 | 9.1 | 1.0 | 2.3 | 0.08 |
| (Fr. 1.4M) | 11.8 | 11.7 | 1.0 | 2.7 | 0.10 |
| (Fr. 2.0M) | 17.3 | 17.0 | 1.0 | 3.4 | 0.15 |

PPS and its fractions, except for fraction Fr. 2.0M, with the highest molecular weight, were analysed by LC-MS. The LC-MS profiles with the identified components are shown in **Fig. 2** with the m/z data and retention times of the components. Some additional minor structures identified are also reported in **Fig. S2**. Overall, the fractions differ from each other principally in the length of oligosaccharides, whereas all the structural groups (e.g. sulfate groups, MGA and acetyl moieties) identified in PPS are homogeneously distributed in each fraction, as confirmed by NMR analysis described below.

**Fig. 2.**
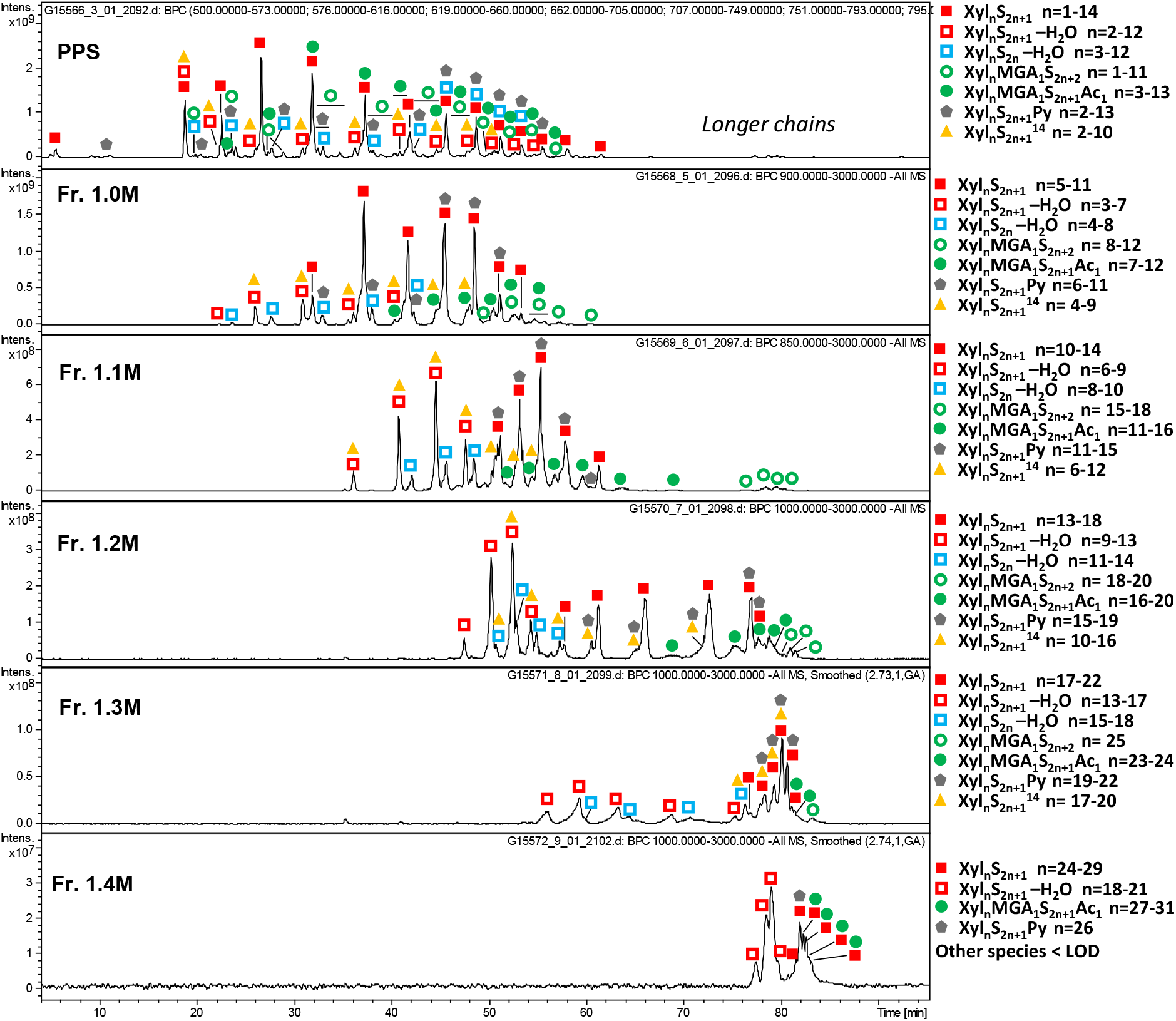
Isothermal calorimetry profiles of PPS and PPS fractions with the identified components. Xyl – xylose repeating unit, MGA – branching 4-O-methyl-glucuronic acid, S – sulfate group, Ac – acetyl group, Py – pyridine moiety.

All the fractions contain linear O-sulfated oligomers Xyl_n_S_2n+1_ with one unsubstituted hydroxyl group at C_1_ of the reducing end (RE) (**Fig. 1A**) as prevalent components (**Fig. 2**). Oligosaccharides with two unsubstituted hydroxyl groups (Xyl_n_S_2n_) are detected only at trace level, confirming essentially the complete sulfation of the internal chains (**Fig. S2**). Minor oligosaccharides with an additional 14Da mass shift Xyl_n_S_2n+1_ generally accompany the most abundant Xyl_n_S_2n+1_ species and show multiple peaks in the LC-MS profile, most likely representing various positional isomers related to the presence of L-rhamnose instead of D-xylose.

For all fractions, ion exchange chromatography-based fractionation led to co-elution of regular Xyl_n_S_2n+1_ oligosaccharides with dehydrated Xyl_n_S_2n+1_-H_2_O oligomers, these latter being present within shorter sequences. The loss of a water molecule from the RE, for Xyl_n_S_2n_-H_2_O, containing a C2-C3 double bond at the RE (**Fig. 1A**), explains why the internal sequences are the same as for Xyl_n_S_2n+1_ oligosaccharides. The co-elution of shorter dehydrated oligomers with Xyl_n_S_2n+1_ suggests that the former are better retained by the ion exchange column, most likely because of their higher polarity. An inverse trend is observed for pyridine containing oligomers. The Xyl_n_S_2n+1_Py envelope is slightly shifted to longer chains (**Fig. 2**) that may be explained by charge compensation between a negatively charged sulfate and positively charged nitrogen atom of a pyridine moiety (**Fig. 1A**). Interestingly, MGA-branched structures, with or without 3-O-acetylation (Xyl_n_MGA1S_2n+1_Ac_1_ and Xyl_n_MGA1S_2n+2_, **Figure 1a**), are also characterized by slightly longer chains than the regular Xyl_n_S_2n+1_ oligomers. Their co-elution might be associated with weaker retention of MGA-branched structures on the anion exchange column due to steric effects. As previously reported, MGA-containing structures often exhibit multiple peaks of positional isomers.^19^

Qualitative and quantitative fingerprints of monosaccharide composition of PPS and its fractions were obtained by ^1^H and HSQC-NMR spectroscopy, according to methods recently described.^19^ (**Fig. 1B** and **Table S1**). The monosaccharide composition was determined by ^1^H NMR (**Table 2**). The data provided quantitative information concerning some structural features of PPS which can be related directly to the xylan source (e.g. MGA content) and its preparation process, such as minor modifications at the RE of PPS chains (e.g. through formation of double bonds, pyridine-derivatives and possible methylation as well as residual acetylation). The RE signal of β-xylose at 5.1 ppm is partially overlaid by that of a xylose residue at 5.08 ppm, the structure of which has yet to be fully elucidated (Xyl*6). The area was calculated using the corresponding α-signal at 5.38 ppm and the α/β ratio (35/65) obtained by 2D NMR. ΔXylred (CHO) was calculated from the integral of the H3 signal at 6.78 ppm, while the level of Xyl3Ac-2MGA was calculated from the acetyl signal at 2.31 ppm. Finally, the degree of acetylation was determined by calculating the ratio of the sum of acetyl groups between 2.35 and 2.1 ppm and the sum of xylose related signals.

**Table 2.** Monosaccharide compositional analysis by ^1^H NMR (molar percentage).

| Residue | PPS | Fr. 1.0M | Fr. 1.1M | Fr. 1.2M | Fr. 1.3M | Fr. 1.4M | Fr. 2.0M |
| --- | --- | --- | --- | --- | --- | --- | --- |
| Xyl+XylNR | 67.5 | 68.9 | 73.6 | 75.5 | 76.4 | 76.9 | 82.4 |
| XylRE | 14.3 | 13.4 | 7.5 | 5.6 | 4.4 | 3.7 | 2.9 |
| Xyl3Ac-2MGA | 2.3 | 2.6 | 2.7 | 2.7 | 2.6 | 2.6 | 1.9 |
| $\Delta\text{Xyl}$ | 0.6 | 0.4 | 0.3 | 0.3 | 0.2 | 0.1 | 0.2 |
| $\Delta\text{Xylred(CHO)}$ | 1.1 | 1.0 | 1.3 | 1.0 | 0.8 | 0.5 | 0.6 |
| Xylα(Py)<br>+Xylβ(Py) | 1.4 | 1.3 | 0.6 | 0.5 | 0.3 | 0.2 | 0.2 |
| MGA-(Xyl3Ac) | 2.2 | 2.2 | 2.5 | 2.7 | 2.6 | 2.5 | 1.9 |
| MGA-(Xyl) | 1.1 | 1.2 | 0.8 | 0.9 | 0.9 | 0.7 | 0.7 |
| MGA*-(Xyl) | 1.0 | 0.8 | 0.3 | 0.3 | 0.2 | 0.2 | 0.6 |
| Degree of<br>acetylation | 4.1 | 4.8 | 5.1 | 5.2 | 5.2 | 5.5 | 4.5 |

The advantage of the quantitative 2D-NMR method^24,25^ consists in the enhancement of signal dispersion compared to the mono-dimensional spectra, which enables better resolution of additional minor signals of PPS. The molar percentages of each residue (**Table S1**) were calculated from the abundance of corresponding cross peak volumes normalized to the sum of all the cross-peak volumes.^19^ Both mono- and bi-dimensional techniques indicated the structural similarity for all PPS fractions, which differ mainly by the quantity of their reducing and non-reducing residues. (**Table 2** and **Table S1**).

### ITC determination of SARS-CoV-2 S1-RBD Binding to PPS

Isothermal titration calorimetry (ITC) is a sensitive technique for detecting biomolecular interactions by measuring the heat absorbed or released upon binding. ITC does not require immobilization or labelling of the partners and can be performed in solution, thereby measuring the affinity of binding molecules in their native state. In addition, ITC experiments provide not only the parameters relating to binding, such as binding constant (K_D_) and stoichiometry (N), but also the enthalpy and entropy contributions of that binding (ΔH, ΔS) providing information concerning the nature and magnitude of the forces driving the binding. To evaluate the possible correlation between the binding affinity and the PPS chain length, ITC experiments were performed to address the interaction between the S1-RBD and PPS fractions with different average chain lengths. ITC experiments were first performed in phosphate buffer solution (PBS) that mimics the pH, osmolarity and ion concentration of human fluids. The ITC profiles of S1-RBD titrated with PPS and its fractions were fitted using two sets of binding site models which indicated a strong primary binding site and a non-specific secondary binding site (**Table S2**). To minimize the non-specific binding, we performed the experiments increasing the salt concentration. At higher ionic strength (500 mM), the added salt led to full screening of the electrostatic interaction between PPS chains and S1-RBD with no evident heat changes in the ITC profiles. Furthermore, an increase of the salt concentration from 140 mM to 200 mM resulted in non-reproducible ITC titrations, despite the protein sample and experimental conditions remained unchanged during repeated measurements. We therefore changed the buffer solution from PBS to HEPES, that is another widely used buffer in biological studies, suitable for ITC experiments due to its low ionization enthalpy. The state and conformational stability of the protein in HEPES buffer was further confirmed by DLS measurements and CD spectra (**Fig. S3**)

ITC thermograms resulting from the titration of S1-RBD in HEPES buffer using the same PPS fractions that were employed in the *in vitro* antiviral activity measurements are shown in Figure 3. The ITC profiles showed exothermic heat of binding that could be fitted using a one-site binding site model (**Table 3**). The binding affinity was found to increase with the increasing length of ligand, as the apparent dissociation constant decreased as ligand molecular weight increased (from a K_D_ of 8.41 x 10^-7^ M to 7.42 x 10^-8^ M). For all fractions, N values were in the range 0.18 to 0.24, indicating that 4–5 molecules of S1-RBD were bound to a single molecule of PPS. A similar binding stoichiometry had already been observed for PF4 and FGF1 which form multimolecular complexes in the presence of heparin.^26,27^ The binding to the S1-RBD resulted in negative binding free energy that became slightly more favourable at increasing ligand molecular weights (−8.29 to −9.73 kcal/mol). All interactions had both favourable enthalpy (−0.96 to −5.78 kcal/mol) and entropy (−3.15 to −7.72 kcal/mol) terms. The negative enthalpic contribution to binding was mainly due to the formation of strong ionic interactions, in addition to hydrogen bonds and van der Waals interactions between the side chains of basic residues on the S1-RBD surface and the sulfate groups of the ligand. The favourable entropic contribution was presumably due to the higher release of water molecules to bulk upon ligand and target interaction.

**Fig. 3.**
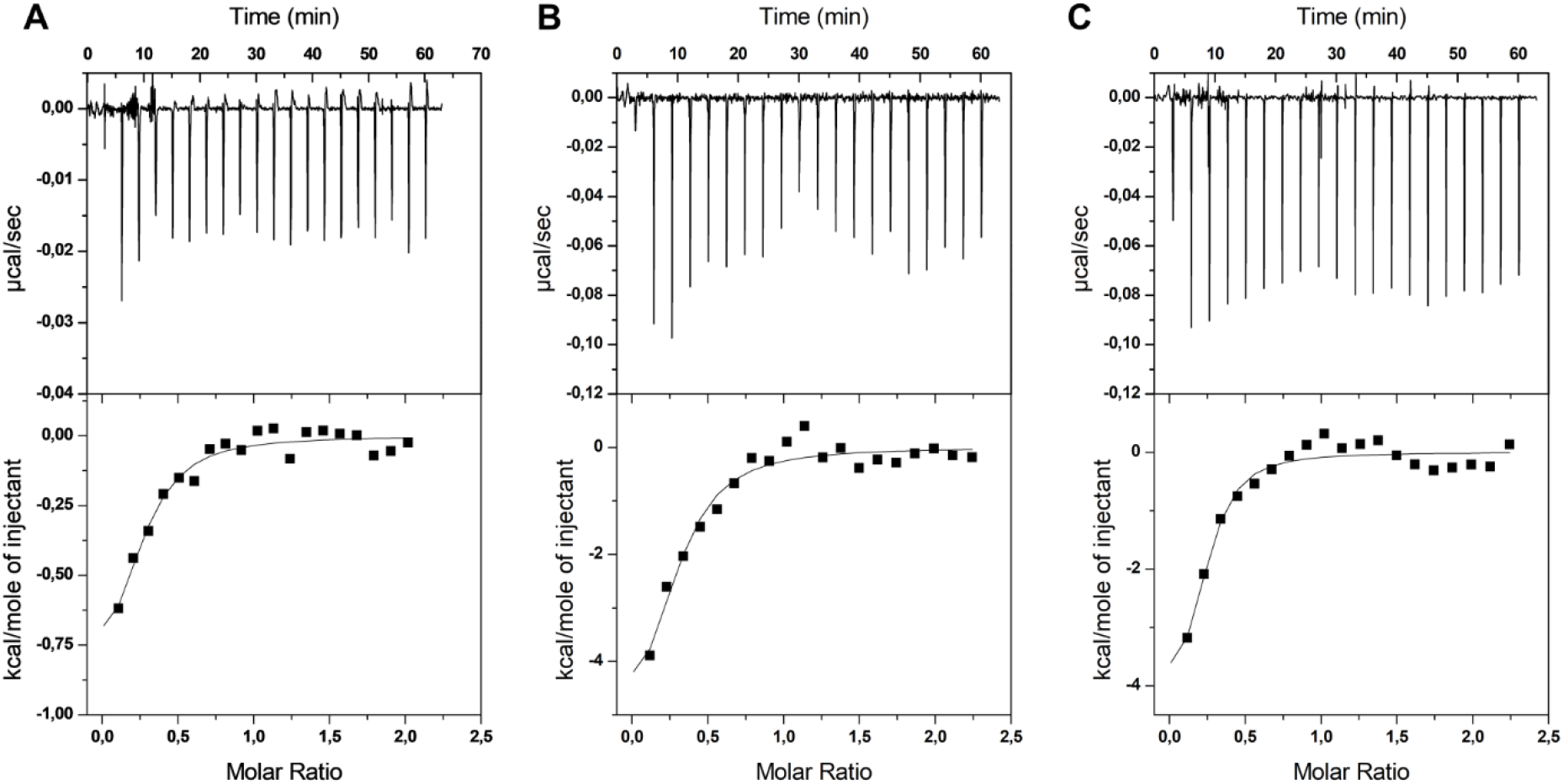
Representative profiles of ITC titration at pH 7.2 of S1-RBD domain with fractions: A) Fr-1.0 M, B) Fr-1.2 M and C) Fr-1.4 M PPS in 20 mM HEPES buffer and 200 mM of salt. The upper panels show the raw data (heat pulse for every injection). The bottom panels represent integrated areas per mole of injected ligand as a function of molar ratio. The solid line is the best fit to the experimental data using a single-site binding model.

**Table 3.** Thermodynamic parameters obtained from ITC titrations of PPS fractions into S1-RBD domain using a single-site binding model. Titration was performed in 20 mM HEPES buffer pH 7.4 and 200mM NaCl. (* 140mM NaCl)

| | MW (KDa) | N (sites) | $K_D$ (M) | $\Delta H$<br>(kcal/mol) | $\Delta G$<br>(kcal/mol) | $-T\Delta S$<br>(kcal/mol) |
| --- | --- | --- | --- | --- | --- | --- |
| PPS* | 5.7 | $0.20 \pm 0.01$ | $6.13 \times 10^{-7} \pm 3.96 \times 10^{-7}$ | $-2.20 \pm 1.45$ | -8.48 | -6.28 |
| Fr-1.0M | 3.7 | $0.24 \pm 0.05$ | $8.41 \times 10^{-7} \pm 5.36 \times 10^{-7}$ | $-0.97 \pm 0.34$ | -8.29 | -7.31 |
| Fr-1.1M | 5.3 | $0.18 \pm 0.11$ | $4.33 \times 10^{-7} \pm 3.48 \times 10^{-7}$ | $-0.96 \pm 0.74$ | -8.68 | -7.72 |
| Fr-1.2M | 7.0 | $0.24 \pm 0.05$ | $2.85 \times 10^{-7} \pm 1.58 \times 10^{-7}$ | $-5.78 \pm 1.80$ | -8.93 | -3.15 |
| Fr-1.3M | 9.2 | $0.23 \pm 0.04$ | $1.84 \times 10^{-7} \pm 1.00 \times 10^{-7}$ | $-4.02 \pm 0.99$ | -9.19 | -5.17 |
| Fr-1.4M | 11.8 | $0.22 \pm 0.04$ | $1.61 \times 10^{-7} \pm 1.07 \times 10^{-7}$ | $-4.64 \pm 1.34$ | -9.27 | -4.63 |
| Fr-2.0M | 17.3 | $0.22 \pm 0.02$ | $7.42 \times 10^{-8} \pm 3.69 \times 10^{-8}$ | $-5.24 \pm 0.70$ | -9.73 | -4.49 |

In contrast with PPS fractions, titration experiments performed using the PPS mixture as ligand showed no interactions at 200 mM NaCl, consistent with a low affinity of binding (K_D_ in the mM range), which could not be detected with ITC measurements (**Fig. S4**). However, calorimetric titrations performed decreasing the salt concentration at 140mM resulted in a binding isotherm with a K_D_ of 6.13 x 10^-7^ ± 3.96 x 10^-7^ M and with a binding that was enthalpically (ΔH= −2.20 kcal/mol) as well as entropically (-TΔS= −6.28 kcal/mol) driven. The resulting binding stoichiometry was similar to that found for the fractions, where 4-5 S1-RBD molecules bind 1 molecule of PPS. To confirm that high salt concentrations screened any electrostatic interactions between PPS and S1-RBD, an ITC titration was performed, lowering the salt concentration to 60mM (**Fig. S4**). The fitted data yielded a lower dissociation constant (K_D_, 1.64 x 10^-7^ + 0.47 x 10^-7^ M), suggesting that electrostatic interactions contribute significantly to the affinity between the PPS and the S1-RBD domain.

### Structural characterization of SARS-CoV-2 S1-RBD upon binding to PPS

Since no significant changes were observed in the hydrodynamic radius of the S1-RBD in complex with the PPS chains, we investigated whether alterations in the S1-RBD structure accompany the binding. Far-UV CD spectroscopy, that is sensitive to conformational changes in proteins upon ligand binding, was employed. Specifically, the CD signal in the far-UV (200–250 nm) provides information concerning changes in the proportions of secondary structural elements (α-helices, β-sheet and unstructured regions) that could take place following ligand binding. The far-UV CD spectra acquired on S1-RBD at increasing ligand concentration reported only slight changes of secondary structure, without highlighting any trend related to increased concentrations of PPS or its fractions (**Table 4**). In agreement with the DLS data, no increase in antiparallel β-sheet content was measured, which can indicate aggregation, nor was an increase in the level of signal amplification required (which would have been indicative of scattering arising from aggregation). These results suggest that the various ligands tested induce broadly similar conformations in the protein and are consistent with the interpretation that no significant aggregation was associated with the binding.

**Table 4.**
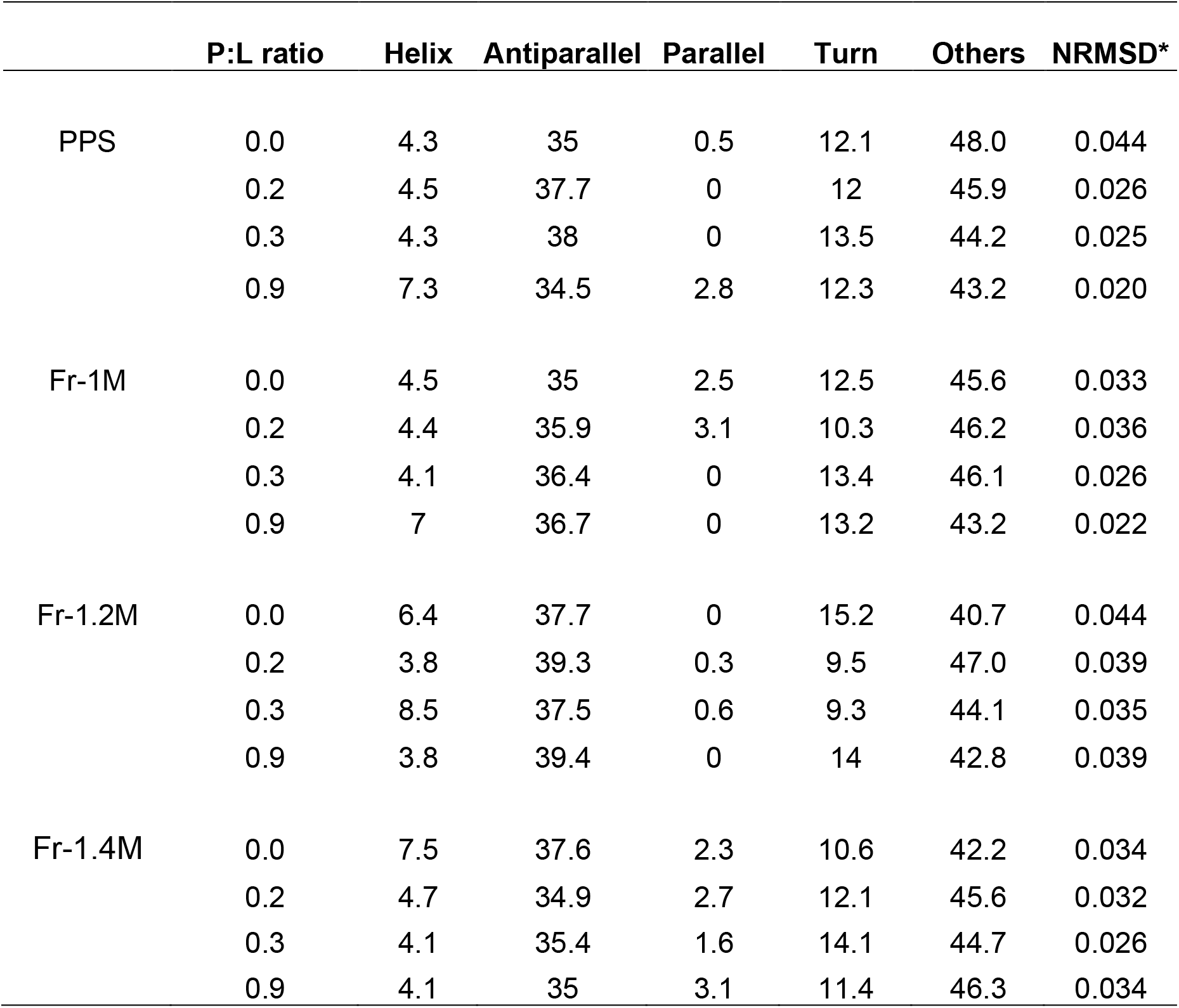
Conformational changes in S1-RBD upon addition of PPS or PPS fractions, determined by far UV-CD.

|  | P:L ratio | Helix | Antiparallel | Parallel | Turn | Others | NRMSD* |
| --- | --- | --- | --- | --- | --- | --- | --- |
| PPS | 0.0 | 4.3 | 35 | 0.5 | 12.1 | 48.0 | 0.044 |
|  | 0.2 | 4.5 | 37.7 | 0 | 12 | 45.9 | 0.026 |
|  | 0.3 | 4.3 | 38 | 0 | 13.5 | 44.2 | 0.025 |
|  | 0.9 | 7.3 | 34.5 | 2.8 | 12.3 | 43.2 | 0.020 |
| Fr-1M | 0.0 | 4.5 | 35 | 2.5 | 12.5 | 45.6 | 0.033 |
|  | 0.2 | 4.4 | 35.9 | 3.1 | 10.3 | 46.2 | 0.036 |
|  | 0.3 | 4.1 | 36.4 | 0 | 13.4 | 46.1 | 0.026 |
|  | 0.9 | 7 | 36.7 | 0 | 13.2 | 43.2 | 0.022 |
| Fr-1.2M | 0.0 | 6.4 | 37.7 | 0 | 15.2 | 40.7 | 0.044 |
|  | 0.2 | 3.8 | 39.3 | 0.3 | 9.5 | 47.0 | 0.039 |
|  | 0.3 | 8.5 | 37.5 | 0.6 | 9.3 | 44.1 | 0.035 |
|  | 0.9 | 3.8 | 39.4 | 0 | 14 | 42.8 | 0.039 |
| Fr-1.4M | 0.0 | 7.5 | 37.6 | 2.3 | 10.6 | 42.2 | 0.034 |
|  | 0.2 | 4.7 | 34.9 | 2.7 | 12.1 | 45.6 | 0.032 |
|  | 0.3 | 4.1 | 35.4 | 1.6 | 14.1 | 44.7 | 0.026 |
|  | 0.9 | 4.1 | 35 | 3.1 | 11.4 | 46.3 | 0.034 |

### PPS-Binding site analysis

The determination of the possible contacts between a 2,3-disulfated polyxylan oligosaccharide and the S1-RBD of the SARS-CoV-2 was performed by molecular docking simulation, in which the hexasaccharide 2,3,4-trisulfo-Xyl-β(1-4)-[2,3-disulfo-Xyl]_5_-OH was docked onto the convex surface of the S1-RBD (receptor). The conformation of 2,3-disulfo-xylan residues was set in the ^1^C_4_ conformation as suggested by ^3^J_H-H_ couplings, nuclear Overhauser effect contacts and molecular modelling analysis (**Fig. S5**). The xylohexasaccharide model was built in two limit conformations, setting all the xylose (Xyl) residues in ^1^C_4_ and ^4^C_1_ chair forms. After energy minimization, the xylohexasaccharide with residues in ^1^C_4_ showed the lowest potential energy (−5318.81 kJ/mol) compared to the ^4^C_1_ conformation (−5281.10 kJ/mol). Interestingly, the ^1^C_4_ xylohexasaccharide allows greater distances between the two sulfate groups located in 2-O- and 3-O- of each Xyl residues (**Fig. S5**). The observed inversion of conformation of the xylose pyranoside ring from ^4^C_1_ to ^1^C_4_, resulting from repulsion between the negatively charged sulfate groups, has also been observed for the internal chain residues of 2-3-disulfo-Xylo-oligosaccharides^28^ and the glucuronic acid residue of chondroitin, following full chemical O-sulfation.^29^

The targeted site on the S1-RBD was identified by the series of amino acids that are characterized by positively charged side chains: R346, N354, R355, R357, K444, R466, which remain accessible from the solvent when the S1-RBD is in the ‘up’ conformation on the trimeric S protein. This region has been previously identified as the preferred molecular recognition site toward glycosaminoglycans.^6^ The docking simulation was set up centring the grid box on the Cα of K356 and taking care to leave suitable space for the ligand to be adjusted on the surface of the S1-RBD, considering all possible relative orientations between the ligand and the receptor that were explored in the automatic search procedure. The docked solutions were clustered (tolerance < 2.0 Å), and the clusters ranked by the predicted binding energy (Autodock 4.2 score function). Clusters of docking solutions were then selected considering the binding energy and the population of the clusters. Analysis of the docking solutions suggested weak specificity in the molecular recognition between the hexasaccharide and the S1-RBD of SARS-CoV-2. In fact, 22 clusters of poses (populated by from 1 to 6 poses) presented binding energies between −3.5 to −1 kcal/mol. Interestingly, the lowest binding energy structures (−3.5 kcal/mol) correspond to two different clusters of docking poses. These two sets of docking solutions are reported in **Fig. 4**, showing contacts between the amino acids of the site I identified on the S1-RBD and the sulfate groups of the hexasaccharide. These solutions are located at the edge of the β-sheet that includes the site I sequence (N354, R355, K356, R357), and corresponds to the shallow groove delimited by the short helix (C336 and R346) from one side and the loop (P463-S469) to the other side (**Fig.4**). In this case both docking solutions present comparable contact distances (2.7-4.0 Å) between the residues of the site I core (N354, R355, K356, R357) and the nearest sulfate groups of the hexasaccharide.

**Fig. 4.**
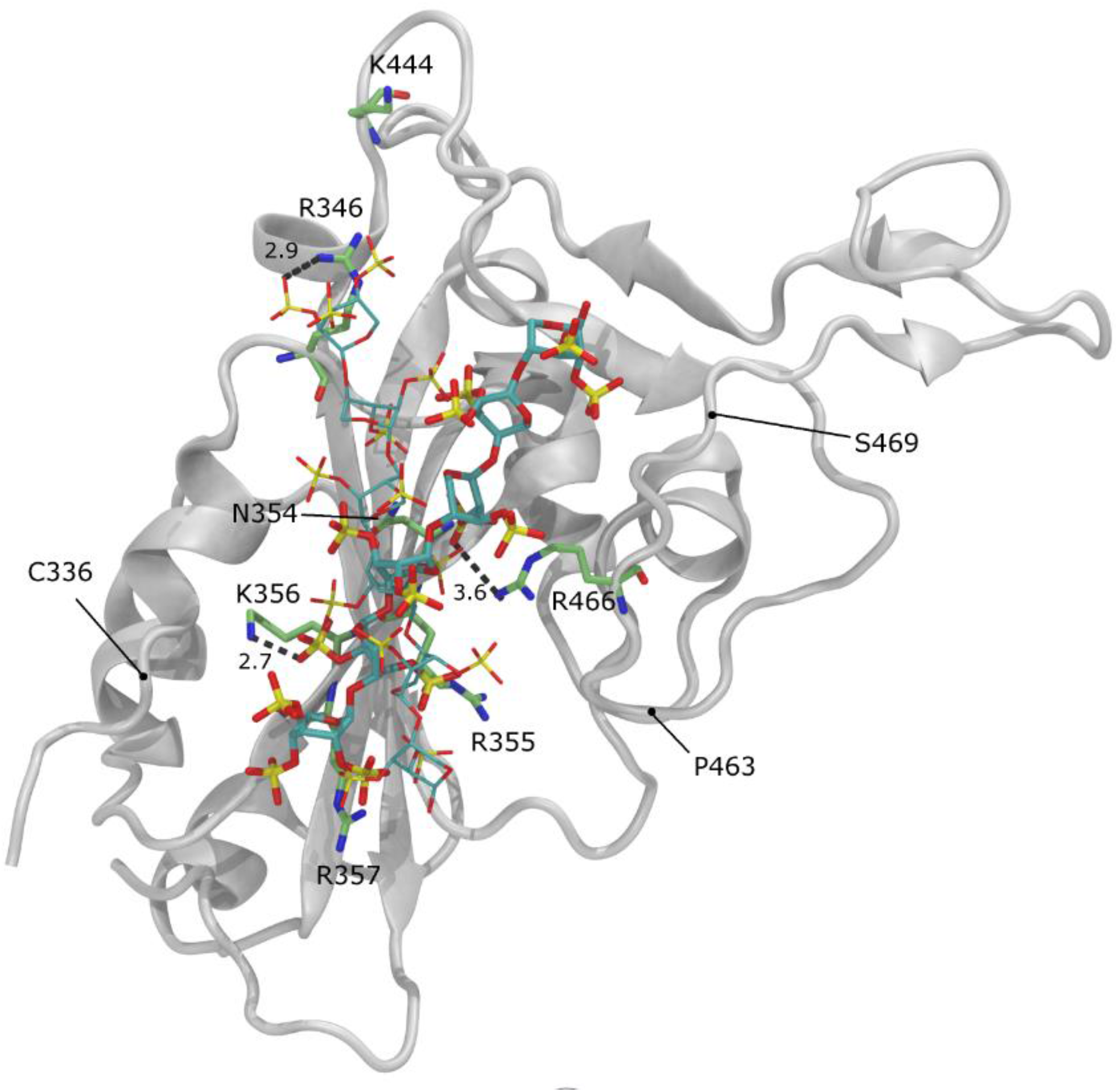
Xylohexasaccharide 2,3,4-trisulfo-Xyl-β(1-4)-[2,3-disulfo-Xyl]_5_-OH docked on the RBD-S1 of the SARS-Cov-2. The representations correspond to the two favorites docking solutions, degenerated in terms of binding energy, but showing different molecular recognition patterns (RMSD > 2.0 Å). Selected distances between the sulfate groups of the ligand and the side chains of selected amino acids are underlined by dashed lines and reported in Å. The RBD-S1 is shown in white ribbon. The amino acids that belong to the site I of the RBD-S1^6^, drawn in green, blue and red tubes, represent carbon, nitrogen and oxygen atoms, respectively. The concave ACE2 binding surface is oriented to the top of the image. The approximate positions of C336, P463 and S469 are underlined to help the reader to localize the helix (C336-R346) and the loop (P463-S469) delimiting the recognition for linear poly-sulfated glycans (see text)

### SARS-CoV-2 Viral Plaque-Forming Assays and S1-RBD Binding

It has been established that the anionic, linear cell surface polysaccharide, HS, is an endogenous receptor for several pathogenic viruses^8,9,11^, including SARS-CoV-2^6,7^ and that heparin, a closely-related, negatively charged polysaccharide with structural similarities to HS, can inhibit this interaction.^6,7,10^ It has been observed for several protein interactions that other sulfated polysaccharides, including those obtained by chemical sulfation of naturally-occurring materials, can emulate heparin and HS, to induce biological effects.^30,31^ The possibility that PPS, which is itself a semi-synthetic product made by the chemical sulfation of a naturally occurring plant xylan, may possess similar inhibitory activities, was therefore investigated.

PPS is an established and approved treatment for cystitis, which can be administered orally, and shows very low toxicity even at high dose (300 mg/day).^32^ It is also reported to exhibit potentially advantageous anti-inflammatory properties.^20^ This alternative potential inhibitor was therefore explored using an established cell-based model of SARS CoV-2 virus invasion and its potency was found to be similar to UFH (**Fig. 5A**). It was found that PPS was able to inhibit viral invasion in experiments involving all three forms of addition; PPS added to cells (cell), to virus (virus) and when added to both before being mixed (virus + cell) as shown in Figure 5a. All three independent experimental approaches highlighted an inhibitory capacity of PPS at least equal to, if not greater than, that of heparin (UFH) on a weight per volume basis.

**Fig. 5.**
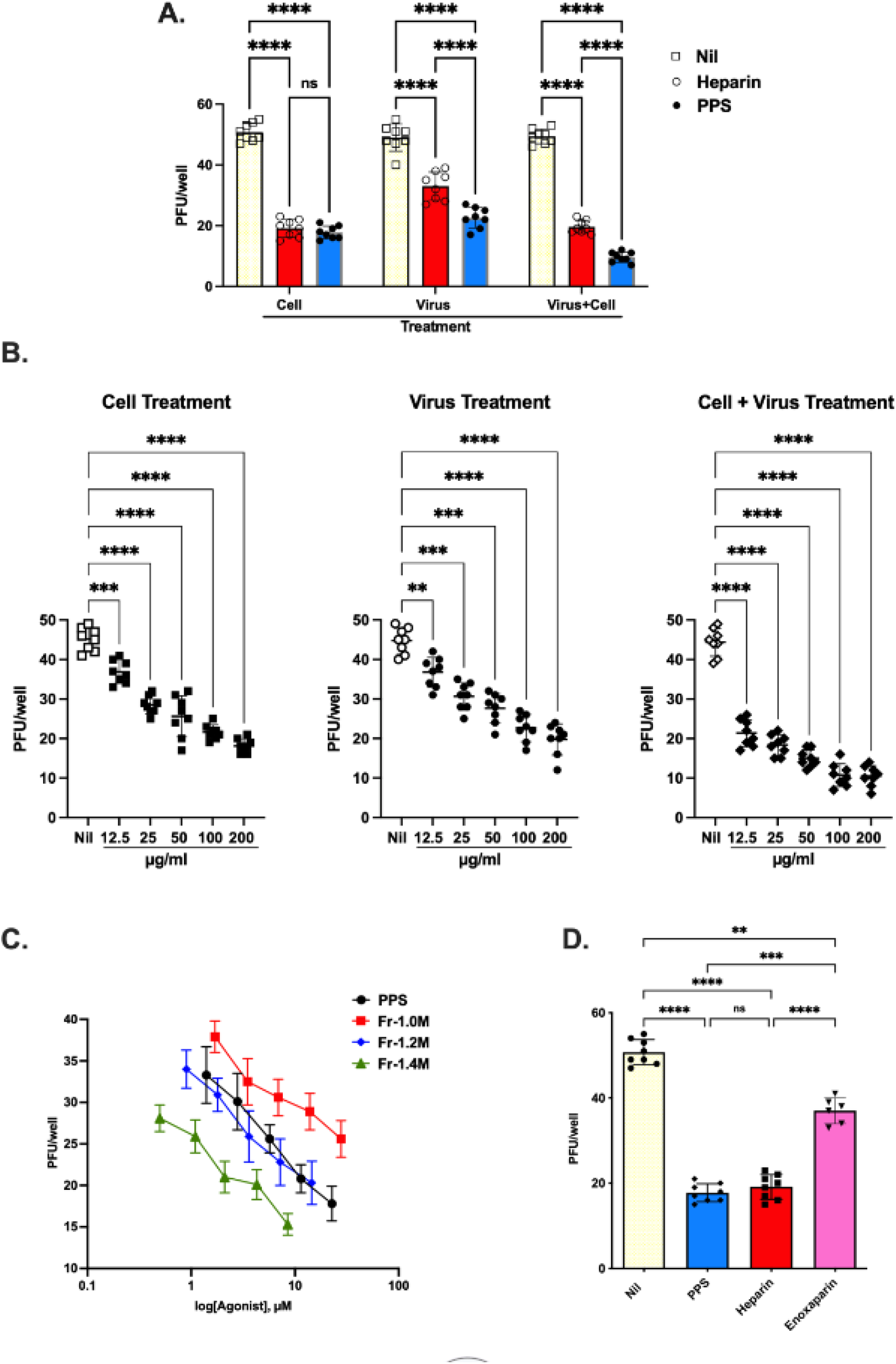
PPS inhibition of SARS-CoV-2 infection. A) Vero cells were treated with PPS and heparin (both at 100 μg/ml) prior to infection with 50 PFU of SARS-CoV-2 or virus-containing supernatant (50 PFU) was treated with the PPS or heparin. In addition, both cells and virus were treated with PPS or heparin as described in materials and methods. B) Dose response when Vero cells were treated with PPS serial dilution prior to infection with 50 PFU of SARS-CoV-2, virus-containing supernatant. (C) Dose response of PPS and PPS fractions-mediated inhibition of SARS-CoV-2 viral invasion of Vero cells. 50 PFU) treated with the PPS serial dilution, both cells and virus were treated with PPS. (D) The PPS-mediated inhibition of SARS-CoV-2 viral invasion of Vero cells (100 μg/mL) compared to heparin and LMWH. One-way ANOVA with Bonferroni correction was used to determine the p values. Nil represents no treatment.

The PPS fractions showed some variation in activity. When compared on a molar basis, a direct MW dependence was observed, the higher molecular weight fraction (Fr-1.4M; Mw 3.7 kDa) being the most active (**Fig. 5C**). However, attributing this with confidence to any particular structural difference is more difficult because the fractions have broadly similar compositions (**Table 2**). The inhibitory efficacy of PPS, UFH and LMWH (enoxaparin) were also compared on a weight per volume basis (**Fig. 5D**). It is clear that PPS is an inhibitor as effective as UFH, when added to cells, on a weight per volume basis (achieving *ca*. 60% inhibition compared to the no inhibitor control) and is more potent than LMWH (which achieved *ca*. 25 % inhibition, (**Fig. 5D**), and which has a closely comparable molecular weight (**Table 1**). Additionally, both PPS and its fractions inhibited viral infection in a dose-dependent manner (in the range 12.5 μg/ml to 200 μg/ml) (**Fig. 5B and 5C,** respectively).

### PPS and fractions exhibit reduced anticoagulant activity compared to heparin

A key advantage of PPS is that it exhibits reduced anticoagulant potential (**Table S3**) which is an important consideration, since high doses may be required in any use as an antiviral agent. Moreover, PPS is less likely to induce bleeding complications. The conventional approach to avoiding the most serious side-effects of prolonged treatment with heparin, such as heparin-induced thrombocytopenia (HIT), is to employ LMWH and a similar strategy could be envisaged in attempting to inhibit SARS-CoV-2.

The anticoagulant activity of PPS and derived fractions was measured as aPTT activity. As expected, a steady increase of anti-coagulation activity with higher molecular weight PPS fractions was observed in the aPTT assay.

## Discussion and Conclusion

Almost two years after the onset of the SARS-CoV-2 pandemic, most countries are still to reach the high levels of vaccination coverage that is necessary to achieve herd immunity. This is compounded by the gradual loss of immunity among those who have already been vaccinated or those who have recovered from an earlier infection and this gradual loss of protection has also been linked to frailty, obesity and deprivation.^33^ The only therapeutic options available currently to treat the early symptoms of coronavirus are injectable antiviral drugs and it was only at the beginning of November 2021 that the first oral antiviral drug (molnupiravir) was approved for use in the UK. There is, therefore, an on-going need to develop additional treatments.

While heparins, the established family of anticoagulant polysaccharides, have shown promise in clinical use^34,35^, reservations remain concerning their direct use as antiviral treatments around potential bleeding complications and the onset of HIT as a side-effect; both of which are associated with high dose and prolonged use. Consequently, alternative agents, which maintain desirable anti-viral properties, while offering reduced anticoagulant and HIT risk, are sought.

Pentosan polysulfate, which is already in use for the relief of bladder pain and discomfort associated with interstitial cystitis has been shown here to possess antiviral properties comparable to those of UFH and stronger than those of LMWH. In contrast to heparin, the lower anticoagulant (aPTT) potency of PPS (10-fold lower compared to heparin) and lack of anti-Xa activity could allow its administration at higher doses thereby enabling its antiviral properties to be exploited.^36,37^ The oligosaccharide fractions of PPS maintain the antiviral properties of the parent intact polysaccharide and this may provide a way of avoiding potential side-effects analogous to HIT, which rely on the formation of immunogenic complexes between platelet factor 4 and HS oligosaccharides of *ca*. 9 kDa or larger.^38^

Both PPS and its fractions bind the S1-RBD efficiently, with Kd values in the μM to nM range, with similar energy changes and both involve interactions that are driven, as expected, primarily by charge-charge interactions (**Table 3**). It is not known whether the detailed interactions of PPS and its fractions with the protein are identical with those of heparin, but docking experiments suggest that the interactions of both heparin and PPS with S1-RBD, occur between the sulfate groups of the xylohexasaccharide and the site I sequence of S1-RBD (N354, R355, K356, R357).^6,7^ An interesting observation is the change from the normally predominant ^4^C_1_ chair form of unsulfated pentosan and xylans^39^, to the ^1^C_4_ conformation for the polysulfated xylose residues. This is presumably the result of repulsion between the charged sulfate groups in position 2 and 3, favouring the lower energy ^1^C_4_ conformation and a greater distance between the two sulfate groups (**Fig. S5**)

The far-UV CD spectra of the S1-RBD alone and the protein in the presence of PPS, while closely similar are, nevertheless, not identical (**Table 4**). While this is broadly consistent with there being no major change of secondary structure, minor changes in protein secondary structure cannot be excluded. The origin of such differences may reside in subtle distinction between the modes of binding of these two polyanions.

It has been noted that, owing largely to our lack of precise understanding at the molecular level of many viral invasion mechanisms, it is highly likely that most current antiviral strategies are not optimised^40^ and this is certainly true for the inhibition of invasion by SARS-CoV-2. It is also noteworthy that both SARS-CoV (CoV-1) and the present SARS CoV-2 have been found to be susceptible to inhibition by anionic polysaccharides^6,8^ and this class of compound warrants more extensive investigation for the inhibition of viral invasion more broadly. Furthermore, PPS and fractions derived from it, provide attractive alternatives to heparin derivatives. They exhibit reduced anticoagulant activity for the inhibition of invasion of susceptible cells by SARS CoV-2 and merit further investigation for potential use as a treatment. While maintaining a better level of inhibitory potential on a weight per volume basis than LMWH and similar to that of UFH, PPS exhibits lower anticoagulant activity. Therefore, PPS should be less prone to bleeding complications at high doses and, by virtue of its distinct structure and the maintenance of anti-viral activity in fractions derived from it and offers an alternative that is likely to be free of HIT or analogous side-effects. The starting material is derived from pentosan, a plant-derived polysaccharide xylan with a well-defined repeating structure^19^ and, even following a chemical sulfation step, it remains highly homogeneous, making it easier to ensure both consistency and to avoid contamination; two problems that have dogged heparin production in recent years.^41,42^

