## Supplemental tables and figures for "Pentosan polysulfate inhibits attachment and infection by SARS-CoV-2 *in vitro*: insights into structural requirements for binding"

### Supplementary

**Table S1** Monosaccharide compositional analysis of PPS and its fractions by HSQC.

| Residue | PPS | Fr. 1.0M | Fr. 1.1M | Fr. 1.2M | Fr. 1.3M | Fr. 1.4M | Fr. 2.0M |
|---|---|---|---|---|---|---|---|
| **Xyl+XylNR** | 71.8 | 71.2 | 77.1 | 79.7 | 82.9 | 82.23 | 86.1 |
| **Xylα** | 4.5 | 4.9 | 2.8 | 2.1 | 1.7 | 1.3 | 0.9 |
| **Xylβ** | 6.7 | 7.3 | 5.3 | 4.5 | 3.8 | 3.6 | 2.7 |
| **Xyl3Ac-2MGA** | 2.1 | 2.3 | 2.3 | 2.3 | 2.0 | 2.1 | 1.4 |
| **ΔXylred** | 1.0 | 0.6 | 0.7 | 0.5 | 0.4 | 0.4 | 0.3 |
| **Xylα(Py** | 0.8 | 0.7 | 0.5 | 0.4 | 0.2 | 0.3 | 0.3 |
| **Xylβ(Py)** | 0.5 | 0.5 | 0.2 | 0.1 | 0.0 | 0.2 | 0.0 |
| **MGA-(Xyl3Ac)+ MGA-(Xyl)** | 4.1 | 4.0 | 3.7 | 3.5 | 3.5 | 3.4 | 2.6 |
| **MGA*-(Xyl)** | 1.0 | 0.9 | 0.4 | 0.3 | 0.3 | 0.2 | 0.0 |
| **Xyl*2** | 0.2 | 0.0 | 0.0 | 0.0 | 0.0 | 0.0 | 0.0 |
| **Xyl*3** | 1.0 | 0.8 | 1.0 | 0.8 | 0.6 | 0.7 | 0.8 |
| **Xyl*4** | 1.3 | 1.3 | 0.8 | 0.6 | 0.4 | 0.4 | 0.4 |
| **Xyl*5** | 0.6 | 0.4 | 0.7 | 0.4 | 0.3 | 0.6 | 0.7 |
| **Xyl*6** | 4.7 | 5.1 | 4.8 | 4.8 | 4.6 | 4.6 | 3.9 |
| **Xyl*7** | 0.7 | 0.8 | 0.6 | 0.5 | 0.5 | 0.5 | 0.3 |
| **MGA tot (from C1)** | 5.1 | 4.8 | 4.1 | 3.8 | 3.7 | 3.6 | 2.6 |
| **MGA tot (from C4)** | 4.8 | 4.8 | 3.7 | 3.5 | 3.5 | 3.5 | 2.7 |
| **3ΔXyl (CHO)** | 1.2 | 0.8 | 1.2 | 1.1 | 0.7 | 0.7 | 0.4 |

**Table S2** Binding affinities and thermodynamic parameters obtained from ITC titrations of PPS fractions into S1-RBD domain in PBS buffer.

| | $N_1$ (sites) | $N_2$ (sites) | $K_{D_1}$ (M) | $K_{D_2}$ (M) | $H_1$ (kcal/mol) | $H_2$ (kcal/mol) | $G_1$ (kcal/mol) | $G_2$ (kcal/mol) | $-T\Delta S_1$ (kcal/mol) | $-T\Delta S_2$ (kcal/mol) |
|---|---|---|---|---|---|---|---|---|---|---|
| PPS | 2.41 ± 0.04 | 0.40 ± 0.03 | 1.11 $x10^{-6}$ ± 3.65$x10^{-8}$ | 3.04 $x10^{-7}$ ± 3.67 $x10^{-8}$ | -4.59 ± 0.98 | -15.7 ± 4.02 | -8.12 | -8.89 | -3.54 | 6.84 |
| Fr-1.0M | 0.80 ± 0.04 | 3.59 ± 0.13 | 1.69 $x10^{-7}$ ± 1.86 $x10^{-8}$ | 1.12 $x10^{-6}$ ± 6.78 $x10^{-9}$ | -4.96 ± 0.58 | -1.41 ± 0.18 | -9.24 | -8.12 | -4.28 | ·6.71 |
| Fr-1.1M | 1.22 ± 0.03 | 5.03 ± 0.06 | 2.93 $x10^{-7}$ ± 4.12 $x10^{-8}$ | 6.27 $x10^{-7}$ ± 4.86 $x10^{-8}$ | -2.50 ± 1.53 | -1.71 ± 0.44 | -8.92 | -8.46 | -6.42 | ·6.75 |
| Fr-1.2M | 1.75 ± 0.09 | 3.67 ± 0.07 | 7.15 $x10^{-7}$ ± 2.93 $x10^{-8}$ | 4.44 $x10^{-7}$ ± 3.53 $x10^{-8}$ | 4.04 ± 3.25 | -5.94 ± 1.42 | -8.39 | -8.67 | -12.4 | ·2.72 |
| Fr-1.3M | 1.61 ± 0.05 | 10.0 ± 0.30 | 2.10 $x10^{-8}$ ± 6.47 $x10^{-9}$ | 1.78x $x10^{-6}$ ± 1.31 $x10^{-8}$ | -3.46 ± 0.49 | -2.75 ± 0.20 | -10.5 | -7.85 | -7.02 | ·5.10 |
| Fr-1.4M | 1.04 ± 0.07 | 10.0 ± 0.70 | 5.69 $x10^{-8}$ ± 1.48$x10^{-8}$ | 1.14x $x10^{-7}$ ± 1.60 $x10^{-8}$ | -1.25 ± 0.52 | -1.23 ± 0.11 | -9.89 | -8.11 | -8.64 | ·6.88 |
| Fr-2.0M | 1.25 ± 0.01 | 9.84 ± 0.05 | 5.85 $x10^{-8}$ ± 3.55 $x10^{-9}$ | 1.39 $x10^{-7}$ ± 3.63 $x10^{-9}$ | -4.06 ± 1.38 | -1.09 ± 0.20 | -9.87 | -9.36 | -5.81 | ·8.26 |

**Table S3** Results of the aPTT assay in seconds. PPS and PPS-fraction were diluted in human pool plasma and tested in the indicated concentrations and blank human pool plasma (0 µg/ml). N/A no result within the linear range of the assay (0 - 250 sec.).

| | 50 µg/mL | 25 µg/mL | 12.5 µg/mL | 6.25 µg/mL | 3.125 µg/mL | 0 µg/mL |
|---|---|---|---|---|---|---|
| PPS | N/A | 180.4 | 81.7 | 50.4 | 39.4 | 31.9 |
| Fr-1.0M | N/A | 121.4 | 63.1 | 45.9 | 38.7 | 31.9 |
| Fr-1.2M | N/A | N/A | 84.3 | 50.6 | 40.1 | 31.9 |
| Fr-1.4M | N/A | N/A | 136.8 | 63.1 | 42.4 | 31.9 |
| Fr-2.0M | N/A | N/A | 185 | 69.7 | 45.0 | 31.9 |

**A**

#### Refractive Index

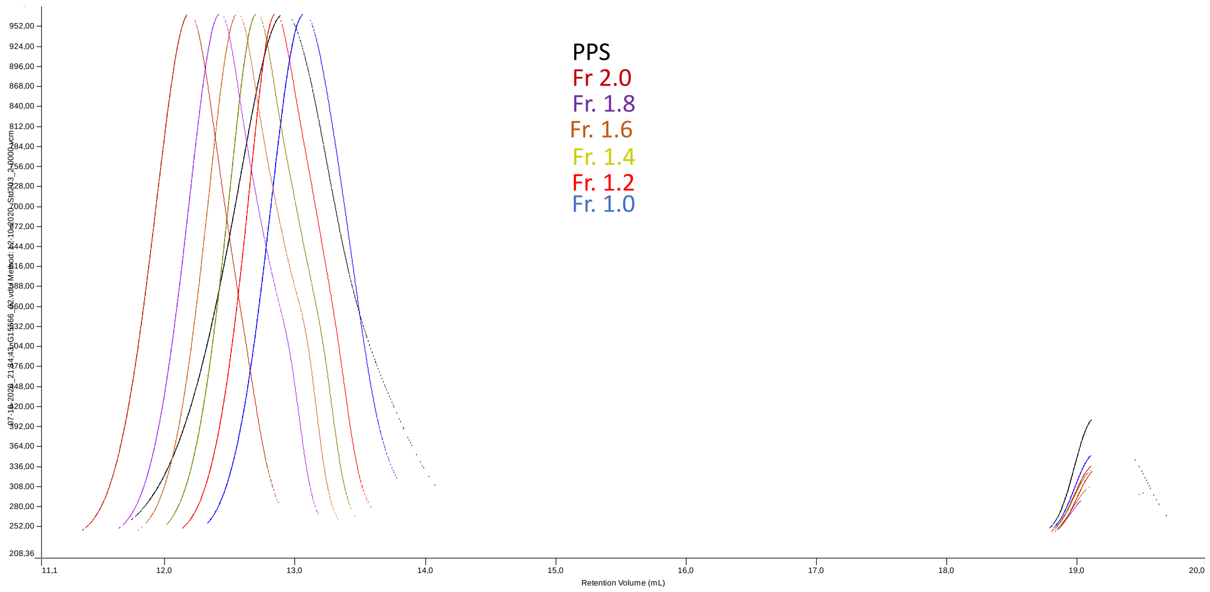


**B**

#### Right Angle light Scattering

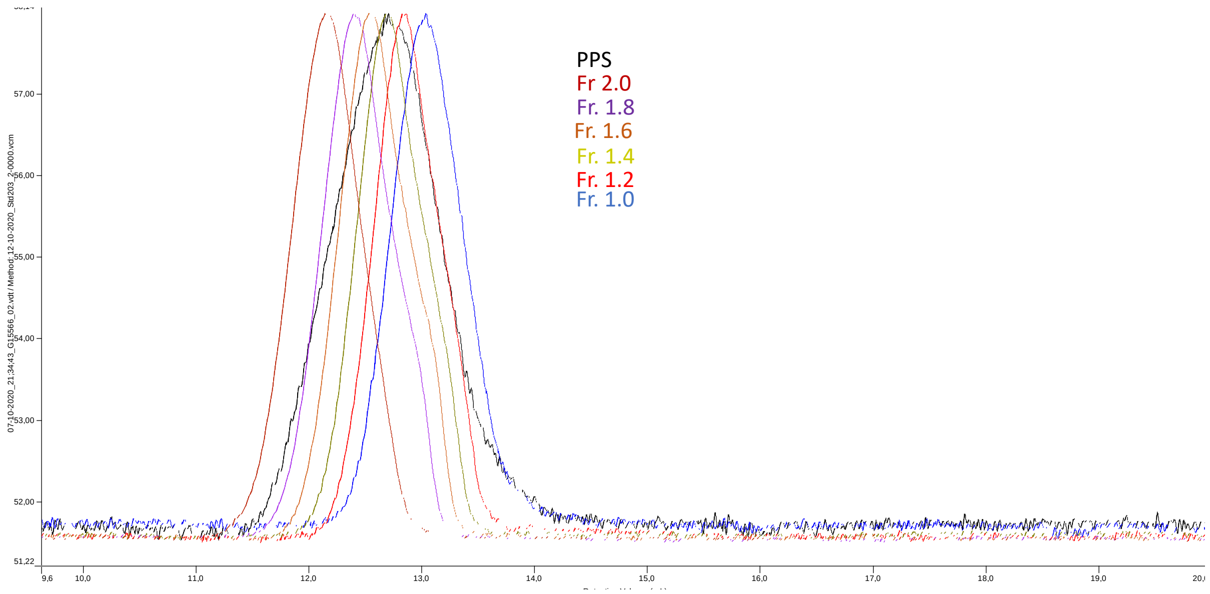


**Figure S1** A) Overlay plots of refractive index vs retention time. B) Overlay plots of right-angle scattering *v.* retention time.

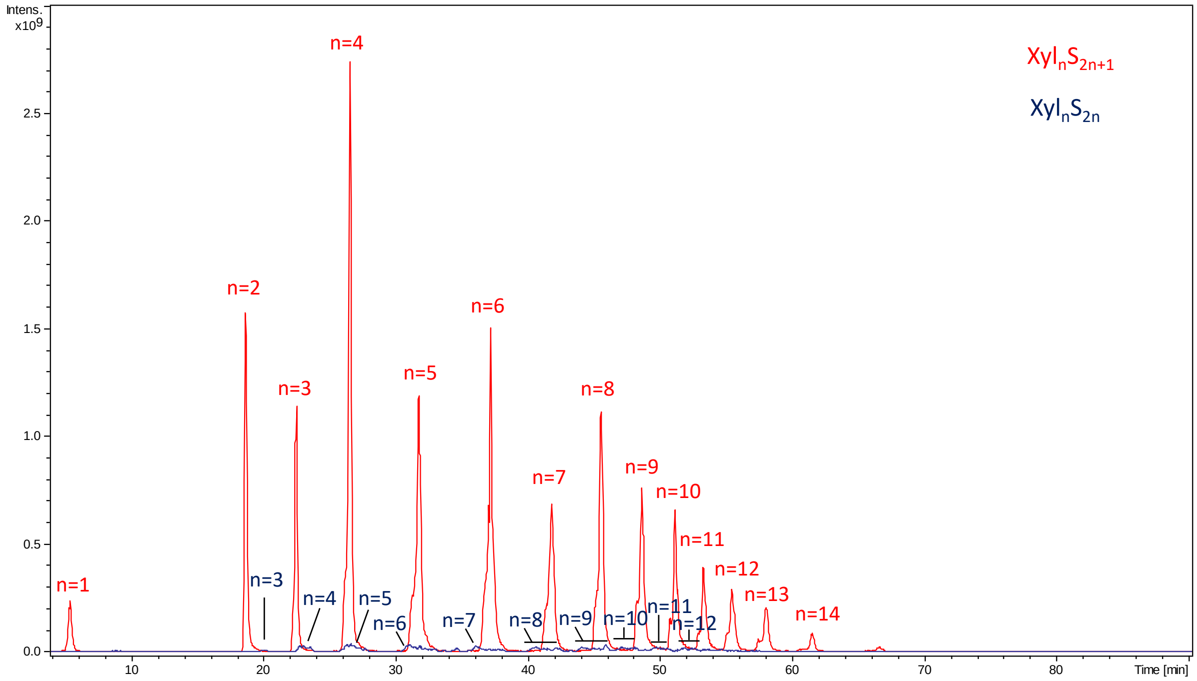


**Figure S2.** Extracted ion chromatograms of most abundant oligosaccharides with fully sulfated internal chains and an unsubstituted hydroxyl group at the RE $Xyl_nS_{2n+1}$ (red) and those with two unsubstituted hydroxyl groups $Xyl_nS_{2n}$ (blue) present only at trace level.

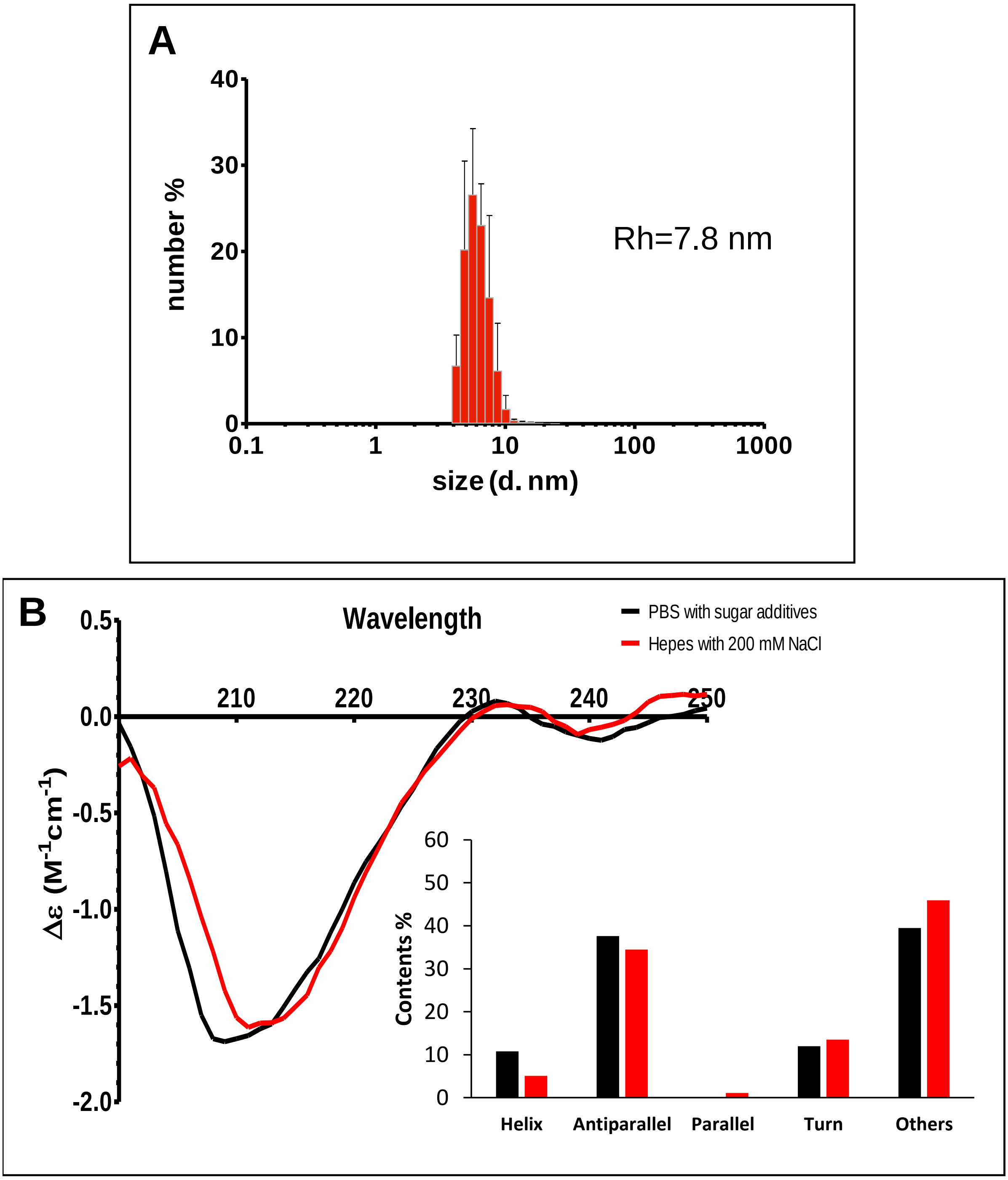


**Fig. S3**. (A) Dynamic light scattering histogram of the molecular size distribution of S1 RDB 1μM in HEPES 20mM, 200 mM NaCl. (B) Circular dichroism spectra (200–250 nm) of SARS-CoV-2 RBD in PBS (black) and in HEPES (red) buffers and corresponding secondary structure content

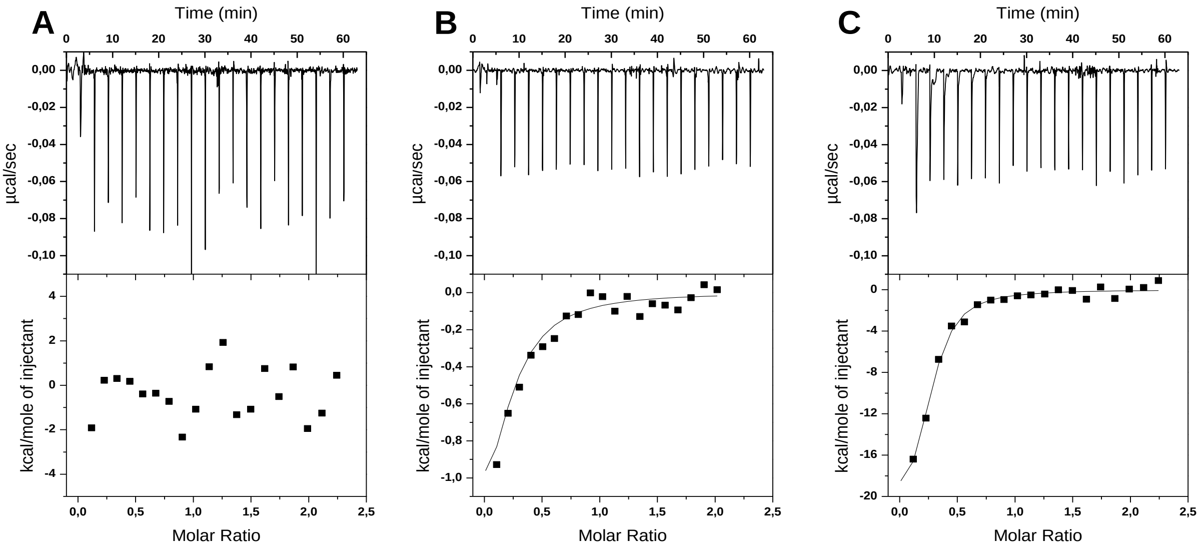


| NaCl (mM) | N (sites) | $K_d$(M) | ΔH (kcal/mol) | ΔG (kcal/mol) | -TΔS (kcal/mol) |
|---|---|---|---|---|---|
| 60 | 0.24 $\pm$ 0.18 | $1.64 \times 10^{-7} \pm 0.47 \times 10^{-7}$ | -23.3 $\pm$ 2.65 | -9.26 | 14.1 |
| 140 | 0.20 $\pm$ 0.01 | $6.13 \times 10^{-7} \pm 3.96 \times 10^{-7}$ | -2.20 $\pm$ 1.45 | -8.48 | -6.28 |
| 200 | - | - | - | - | - |

**Fig. S4**. Interaction of PPS mixture with S1-RBD domain investigated by ITC at different salt concentrations. ITC profiles of S1-RBD titrated with PPS in HEPES buffer pH 7.2 and (A) 200 mM (B) 140mM and (C) 60 mM salt concentration.

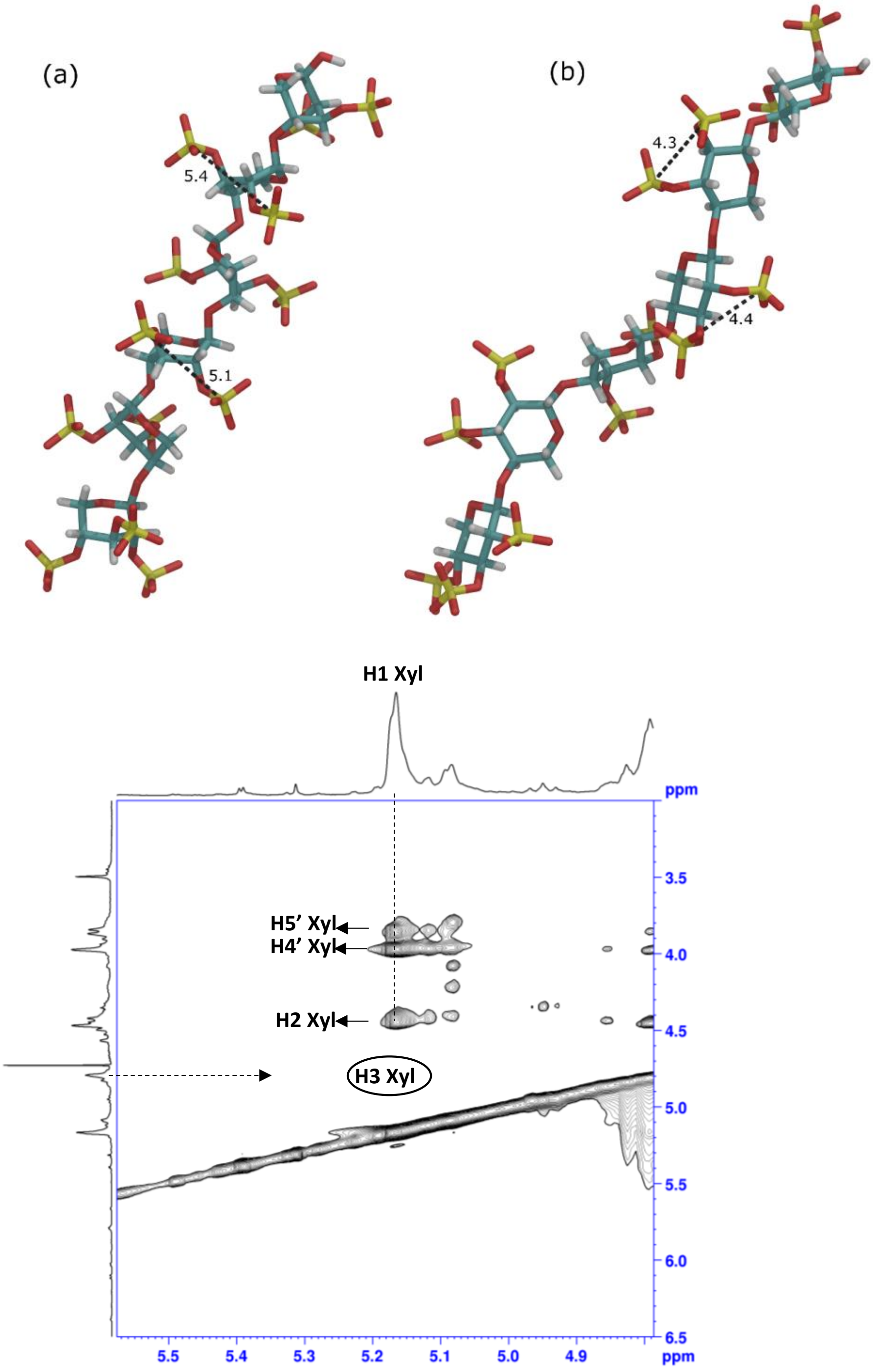


**Fig. S5** Representation of the xylohexasaccharide structure: 2,3,4-trisulfo-Xyl-β(1-4)-[2,3-disulfo-Xyl]5-OH upon energy minimization (see Materials and Methods) in $^{1}C_{4}$ (a) and $^{4}C_{1}$ (b) conformations. The xylohexasaccharide in $^{1}C_{4}$ shows the lowest potential energy and the greater distances between sulfate groups whitin each Xyl residue. (c) 2D-NOESY spectrum of PPS confirming the absence of H1-H3 correlation, characteristic of the $^{4}C_{1}$ conformation and not observable in the $^{1}C_{4}$ form.
